# Host space, not energy or symbiont size, constrains feather mite abundance across passerine bird species

**DOI:** 10.1101/2023.02.03.526976

**Authors:** María del Mar Labrador, David Serrano, Jorge Doña, Eduardo Aguilera, José L. Arroyo, Francisco Atiénzar, Emilio Barba, Ana Bermejo, Guillermo Blanco, Antoni Borràs, Juan A. Calleja, José L. Cantó, Verónica Cortés, Javier De la Puente, Diana De Palacio, Sofía Fernández-González, Jordi Figuerola, Óscar Frías, Benito Fuertes-Marcos, László Z. Garamszegi, Óscar Gordo, Míriam Gurpegui, István Kovács, José L. Martínez, Leandro Meléndez, Alexandre Mestre, Anders P. Møller, Juan S. Monrós, Rubén Moreno-Opo, Carlos Navarro, Péter L. Pap, Javier Pérez-Tris, Rubén Piculo, Carlos Ponce, Heather Proctor, Rubén Rodríguez, Ángel Sallent, Juan Carlos Senar, José L. Tella, Csongor I. Vágási, Matthias Vögeli, Roger Jovani

**Author notes:** Correspondence María del Mar Labrador and Roger Jovani. Doñana Biological Station (CSIC). Avda. Américo Vespucio 26, 41092, Seville, Spain.

## Abstract

Comprehending symbiont abundance among host species is a major ecological endeavour, and the metabolic theory of ecology has been proposed to understand what constraints symbiont populations. We parameterized metabolic theory equations to predict how bird species’ body size and the body size of their feather mites relate to mite abundance according to four potential energy (microbial abundance, uropygial gland size) and space constraints (wing area, number of feather barbs). Predictions were compared with the empirical scaling of feather mite abundance from 26,604 birds of 106 passerine species, using phylogenetic modelling and quantile regression. Feather mite populations were strongly constrained by host space (number of feather barbs) and not energy. Moreover, feather mite species’ body size was unrelated to their abundance or to the body size of their host species. We discuss the implications of our results for our understanding of the bird-feather mite system and for symbiont abundance in general.

## Introduction

A central scope in ecology is to describe abundance patterns, to comprehend the processes that underlay these patterns, and to understand their ecological consequences. These questions have been mainly studied in free-living organisms, while symbiont abundance patterns have received less attention (Cunning & Baker 2004; Dobson et al., 2008). Symbionts (including mutualists, commensals, and parasites) are the most ubiquitous, abundant, and diverse organisms on Earth (Morand 2015; Larsen et al., 2017). They are key components of ecosystems and influence nutrient cycles, food webs, energy flows, and community structure (Hatcher et al., 2012), and their abundance can shape individual host performance and the evolution of host species (Poulin & George-Nascimento 2007). Indeed, the abundance of a given symbiont in or on a given host may determine the nature of the host–symbiont interaction (Bronstein 1994; Holland et al., 2002), with the potential to shift the nature of this relationship between mutualism and parasitism (Hopkins et al., 2017).

Studies on symbiont abundance have mainly focused on parasites rather than on non-parasitic symbionts, and on understanding differences in symbiont abundance among members of a single host species rather than interspecific differences among host species (Turgeon et al., 2018; Mennerat et al., 2021). At the interspecific scale, several studies have found support for Harrison’s Rule which postulates that there is a positive covariation between host size and symbiont size. In contrast, when considering symbiont abundance instead of symbiont size, mixed results have been found for its correlation with the body size of either the hosts or the symbionts (Rózsa 1997a, b; Poulin 1999; Clayton & Walther 2001; Presley & Willig 2008; Krasnov et al., 2013; Galloway & Lamb 2017; Surkova et al., 2018; Lamb & Galloway 2019). At macro-evolutionary scale, host body size largely explained the variation in feather lice effective population size, which is expected to positively correlate with symbiont abundance (Doña & Johnson 2022). Overall, we are still far from understanding why some host species harbour many symbiont individuals of a given taxon, while others carry only a few.

The study of the scaling of symbiont abundance with host body size is an underexplored approach to understand symbiont abundance (Morand & Poulin 2002; George-Nascimento et al., 2004; Poulin & George-Nascimento 2007; Hechinger 2013). Hechinger (2013) developed a hypothesis-driven quantitative framework based on the metabolic theory of ecology (*sensu* Brown et al., 2004) to disentangle how host and symbiont traits shape symbiont abundance across host species. This framework tries to explain symbiont abundance in different hosts through the comparison of theoretical vs. empirical scaling exponents of host and symbiont body size according to energy (e.g. blood or secretions) and space (e.g. surface) provided by the host and according to the metabolic rate and space use of symbionts (see below). Hechinger et al. (2019) used this approach to investigate the relationship between host body size and the abundance of ectosymbiotic mites and lice of 263 bird individuals of 42 species. Their results indicated that the numbers of mites and lice were limited by access to host energy and not by space. However, Hechinger et al. (2019) did not distinguish among ectosymbionts with different diets, e.g. blood-feeding mites were equivalent to non-parasitic mites provided that mite body sizes were similar. Here, we implemented Hechinger’s (2013) framework by analysing an unprecedently large dataset and parametrizing scaling equations using current knowledge of the biology of a particular host–symbiont system: vane-dwelling feather mites (Acariformes: Astigmata: Analgoidea and Pterolichoidea) from European passerine bird species.

Feather mites are ectosymbionts found on almost all birds (Walter & Proctor 2013). Their entire life cycle is spent on their living hosts, mainly on the wing and tail flight feathers, where they are usually queuing between the feather barbs (i.e., the primary branches of the feather rachis; Figure 1) or next to the rachis (Kelso & Nice 1963; Choe & Kim 1989; Yamasaki et al., 2018). They are often said to feed on the preen gland secretions and organic material trapped in them (Dubinin 1951; OConnor 1982; Proctor 2003; Walter & Proctor 2013; Galván et al., 2008). Still, other evidence suggests a lower relevance of preen waxes as food resources (Pap et al., 2010). Algae are also potential food resources for mites (Blanco et al., 2001). However, Doña et al. (2019) studied the gut contents of a large sample of mites using microscopy and DNA metabarcoding, and found that bacteria and fungi were the main food resources for feather mites, while algae and plant materials were rather anecdotic, and bird tissues such as blood or skin were not found.

**Figure 1:**
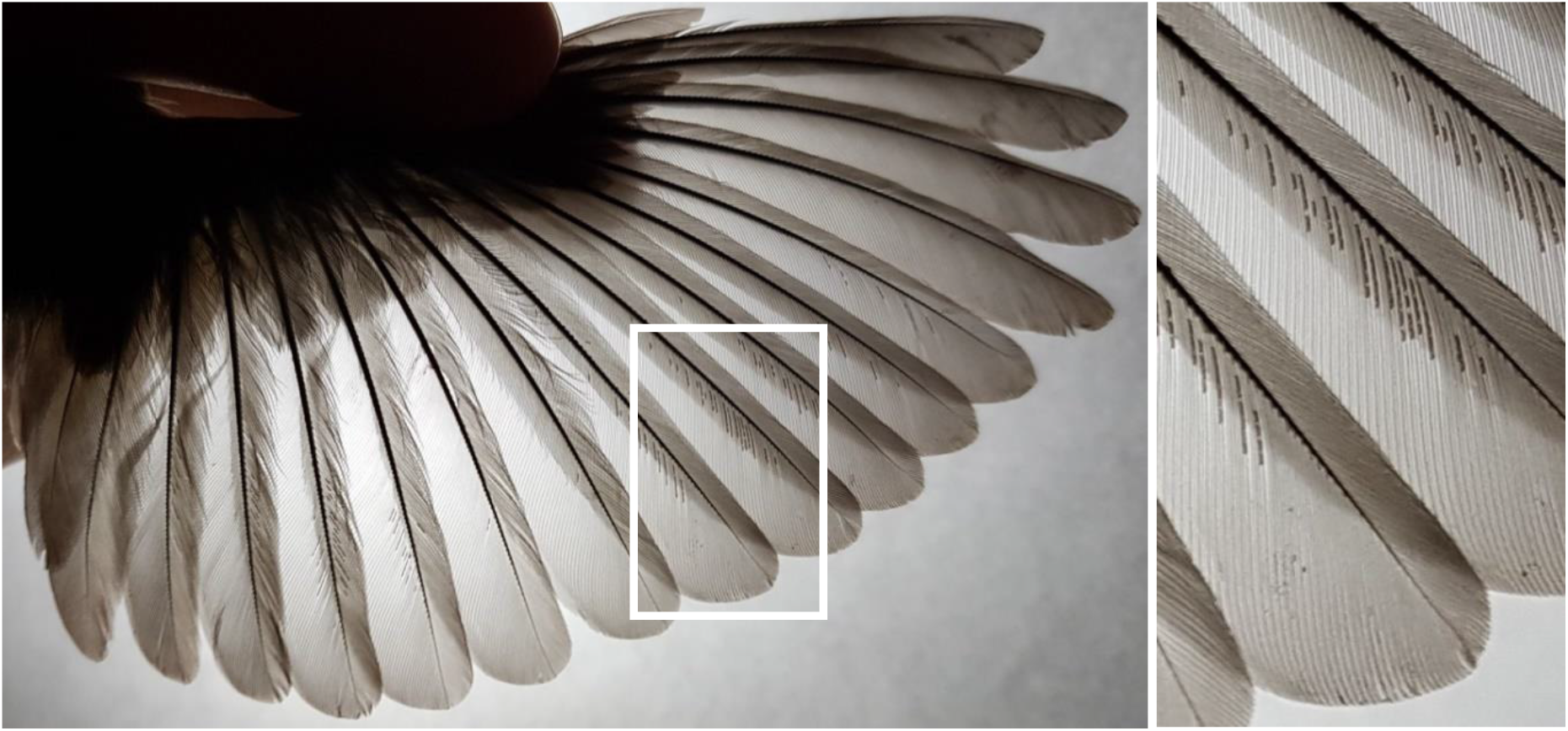
Feather mites (*Proctophyllodes sylviae*) on the wing of a *Sylvia atricapilla*. Note their strong aggregation in certain feathers along the wing and some sections within those feathers, and their queuing along feather barbs.

Bird species strongly differ in feather mite abundance even when accounting for intraspecific variance between localities (Díaz-Real et al., 2014). For instance, species such as *Phylloscopus collybita* and *Periparus ater* consistently have very few feather mites on their wings, while similar-sized *Aegithalos caudatus* and *Acrocephalus melanopogon* often have hundreds of feather mites (Díaz-Real et al., 2014). Interspecific differences in feather mite abundance are partly explained by the ecology and morphology of bird species, but a large proportion of the variance remains unexplained after controlling for these traits (Galván et al., 2008; authors’ unpublished data). To date, only one interspecific study has related bird body size to feather mite abundance (Rózsa 1997b). This study found a positive correlation, albeit based on a relatively small number of host species (N = 17), small number of host individuals within species (range of 3–138), and without quantitatively addressing the underlying mechanisms generating the positive relationship between bird size and feather mite abundance.

In this study, we apply Hechinger’s (2013) quantitative framework to disentangle hosts’ energy and space constraints explaining differences in feather mite abundance across passerine bird species. Here we follow Hechinger’s (2013) use of the term size to refer to the body mass of hosts and symbionts. According to Hechinger (2013), the metabolic theory of ecology predicts that if energy provided by the host (*h*) imposes an effective ceiling to the growth of symbiont (*s*) populations, the maximal or carrying-capacity abundance (but also mean abundance) of the symbiont in a given host individual (*N*_s_) will scale with host body size (*M*_h_) and symbiont size (*M*_s_) as

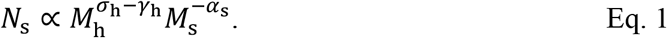

−*γ*_h_ is the scaling exponent for host mass-specific metabolic rate and equals to *α* − 1, where *α* is the scaling exponent for whole-organism metabolic rate to body size (~3/4 across multicellular species; Hechinger 2013). Thus, −*γ*_h_ = −1/4, and −*α*_s_ = −3/4. *σ*_h_ is the spatial exponent for host body size, and is related to the host body part that is metabolically relevant for the studied symbionts (i.e., the host body part that provides the food resources to the symbionts; Hechinger 2013; Hechinger et al., 2019). Current knowledge points to two main energy (food) resources for feather mites, and thus, there are two *σ*_h_ potential values in our study:

1. waxes produced by the uropygial gland that birds spread on feathers (Galván et al., 2008; Doña et al., 2019). We used data on uropygial gland size (see below) to parametrize *σ*_h_ in Eq. 1, given that uropygial gland size is positively correlated, at least within bird species, with the amount of waxes produced (Møller et al., 2009; Pap et al., 2010).
2. organic matter (mainly fungi and bacteria) available on feathers’ surface (Dubinin 1951; Doña et al., 2019; Labrador et al., 2022). This organic matter is not produced by the host, and thus there is not a host body part that is metabolically responsible for its production. We parametrized this alternative *σ*_h_ with data on how the abundance of fungal and bacterial DNA (microbial abundance hereafter) on the wings of passerine bird species scales with bird species body size.

Space provided by the host can also impose an effective ceiling on symbiont populations, and then the maximal or carrying-capacity symbiont abundance in a given host individual would scale with host and symbiont body size as

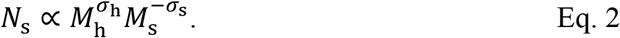

Here, *σ*_h_ indicates how the host body portion that the symbiont inhabits scales to host body size (Hechinger 2013; Hechinger et al., 2019). Theoretical *σ*_h_ values are 1 when the studied symbionts use the host volumetrically, or 2/3 if symbionts inhabit the host surface. Ideally, though, *σ*_h_ should be calculated empirically for each particular study system (Hechinger 2013). We hypothesized that feather mite infracommunities (all of the mite infrapopulations within a single host; Bush et al., 1997) could be spatially constrained by wing area, which is the largest scale habitat for these mites. Alternatively, feather mites could be constrained by the number of feather barbs on the wing because they (except the genus *Trouessartia*) live in the corridors between feather barbs in the ventral side of feathers (Figure 1; Mironov 2022). Moreover, *Trouessartia* spp., despite living on the dorsal surface of feathers (where there are not such well-defined corridors), they also queue along feather barbs (Figure 1 in Mironov & González-Acuña 2013; authors’ personal observation). Thus, we studied the scaling of wing area and the number of barbs to bird species body size to parameterize *σ*_h_ in Eq. 2. Similarly, −*σ*_s_ is the relevant aspect of symbiont bodies that determines their spatial packing on host bodies. Given that mites align in a single row along feather barbs, feather mite length would be the most relevant aspect, and thus we parametrized −*σ*_s_ as −1/3 because this is how mite length scales to mite body size (in μg) (Supporting Information).

In sum we used empirical data to complete the parametrization of Eqs. 1 and 2, and then compared predicted scaling exponents with the empirical exponents obtained by phylogenetic generalized least squares regressions and quantile regressions for the abundance of feather mites across bird species, following Hechinger (2013). We show, using a large dataset on feather mite abundance, how a biologically-informed parametrization of the metabolic theory of ecology proposed by Hechinger (2013) is a powerful approach to understanding why symbiont abundance differs between host species.

## Materials and Methods

### Feather mite morphometric data

Body size in Hechinger’s (2013) equations (*M*_h_, *M*_s_) refers to host and symbiont species’ body masses. Given that these data were available for only one of the mite species studied here, we calculated them from feather mite species’ biometry following the equation provided by Edwards (1967) (Supporting Information). To do so, we gathered data from adult female morphology because they are typically the largest (e.g. Atyeo & Braasch 1966; Santana 1976) and more abundant life stage (e.g. Muzaffar & Jones 2005; Marčanova & Janiga 2021). Feather mites ranged from 394 μm and 0.989 μg for *Scutulanyssus nuntiaeveris* [Berlese] to 1,121 μm and 22.85 μg for *Joubertophyllodes modularis* [Berlese]. Then, to obtain a reliable measure of the mean *M*_s_ on each bird species, we calculated the weighted mean body size (in μg) of the feather mite species reported for each bird species. The weighted mean was calculated using the number of records reported by Doña et al. (2016) for each mite species in each bird species, using only the most reliable bird–mite associations (i.e., those with quality score = 2; see Doña et al., 2016 for more details).

### Feather mite abundance data

Data were obtained from *FeatherMites*, the largest dataset available on feather mite abundances (see Díaz-Real et al., 2014 for details), where, for each bird individual, the total number of vane-dwelling feather mites was counted (i.e., without differentiating between mite species) on the 19 flight feathers (10 primaries, six secondaries, and three tertials) of one wing. Because we aimed to understand the mechanisms setting the upper limit for feather mite abundance, birds without feather mites were not included in the analyses. Therefore, according to parasitological terminology, we analysed feather mite intensity (or infracommunity size; Bush et al., 1997), i.e., the number of feather mites counted in each individual bird with at least one mite, but we use the term ‘abundance’ hereafter due to its general use in the ecology literature. Since we could not find data on the morphology of certain feather mite species in our dataset, some bird species were not included in the analyses, leading to a final dataset of 26,604 individual birds from 106 passerine species.

Given the non-normal frequency distribution of feather mite abundance (Díaz-Real et al., 2014), we used quantiles of mite counts at regular intervals from the 5^th^ (Q5) to the 95^th^ quantile (Q95) to characterize feather mite abundance in each bird species. Special relevance was given to Q95 as the best surrogate of the carrying capacity of feather mite abundance of each bird species, following Hechinger et al. (2019).

### Microbial abundance data

We used microbial abundance data from a recent study where the amount of fungi and bacteria DNA available on feathers’ surfaces was quantified by qPCR (Labrador et al., 2021). This is justified not only by current knowledge on feather mites’ diet (see above), but because Labrador et al. (2022) found that feather mites also occur on wing flight feathers during the night, when they forage. In brief, microbial DNA was extracted and amplified from the second secondary feather of the right wing of 133 individuals of 22 species. The amount of fungal and bacteria DNA were positively correlated at the individual bird and bird species levels (Labrador et al., 2021). Hence, here we combined fungal and bacterial values for each individual bird, and then calculated the mean microbial DNA abundance for each bird species. This value was used as a rough estimate of the microbial food resources available for their feather mites.

### Bird morphology data

Three morphological traits for the studied bird species were retrieved from the literature: body size (in g), wing area, and uropygial gland size (see Supporting Information for details). Moreover, the number of feather barbs was calculated for each bird species combining original data on feather lengths for 40,346 birds (sample size: mean = 917, min-max = 1-9,506 birds per species) captured from 1994 to 2015 at the Manecorro Ringing Station (Doñana National Park, SW Spain), and feather barb density reported in the literature (see Supporting Information for details, and Figure 2 for the number of bird species for each morphological variable).

**Figure 2:**
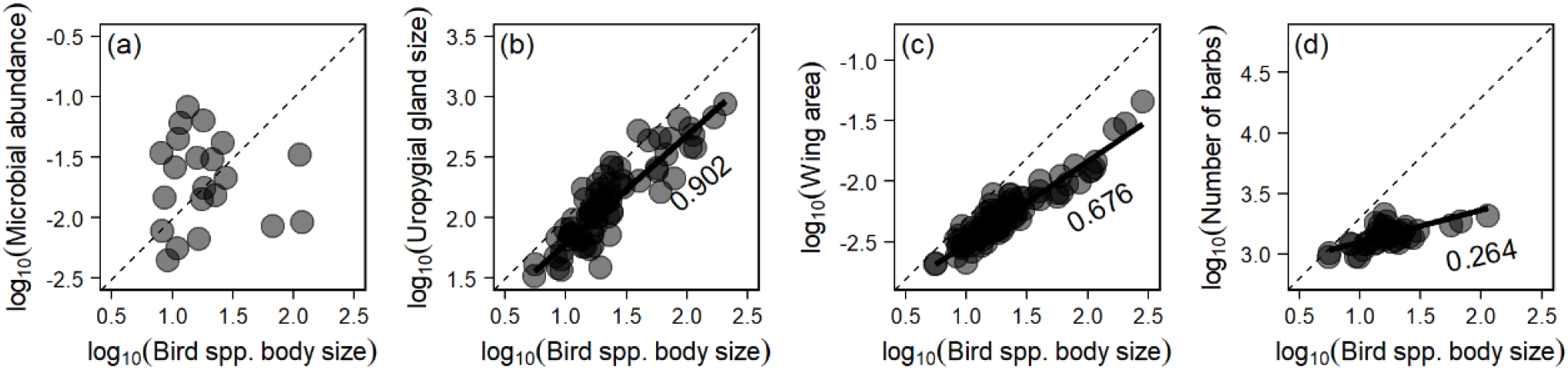
Relationships between potential energetic (microbial abundance N = 21 species, uropygial gland size N = 76) and spatial (wing area N = 88, or number of barbs N = 44) constraints against bird species body size (in g). Dashed lines show slope = 1. Only slopes that departed from 0 (p-value < 0.05) are drawn (black line) and its estimated value is shown.

### Statistical analyses

Phylogenetic generalized least squares regressions (PGLS; Symonds & Blomberg 2014) were performed to retrieve (from the slope of the log-log regressions; following Hechinger 2013) the scaling exponents between bird species’ body size (log_10_ transformed) and the four variables (log_10_ transformed) hypothesized to constrain feather mite infracommunity sizes: wing feather microorganism amount, uropygial gland size, wing area, and number of barbs of primary feathers. One multivariable PGLS regression for each feather mite abundance quantile (dependent variable; log_10_ transformed) was used to calculate how it scaled with bird and mite body sizes (independent variables; log_10_ transformed). PGLS regression was also used to study the relationship between bird size and the weighted mean body size of their feather mites. We used the *gls* function of the *caper* R package (Orme et al., 2012) to perform the PGLS regressions, which ensure the statistical independence of our samples, correcting the model estimates by the phylogenetic relatedness of the studied species. We obtained information on the phylogenetic relationship among bird species by downloading a distribution of 1,000 trees from BirdTree (Jetz et al., 2012, http://birdtree.org) using the Hackett backbone tree (only sequenced species; Hackett et al., 2008). Then, following Rubolini et al. (2015), trees were summarized by computing a single 50% majority-rule consensus tree in SumTrees v 4.5.1 in DendroPy (Sukumaran & Holder 2010, 2015).

In each PGLS model, we allowed the phylogenetic signal in the residuals (i.e., Pagel’s lambda, λ) to be optimized towards its maximum likelihood value (Symonds & Blomberg 2014). These models were also weighted by the sample size (log_10_ transformed) of each bird species to incorporate the higher uncertainty associated with feather mite abundance data from host species with smaller sample sizes.

To further study the factors constraining feather mite infracommunities, we used a multivariable quantile regression analysis on the log_10_(Q95) of feather mite abundance against log_10_(bird body size) and log_10_(feather mite body size) as independent variables (Koenker & Basset 1978; Cade & Noon 2003). We were especially interested in the quantile regressions at the largest τ values because these would reflect the maximum feather mite abundance that bird species can harbour, considering their body size and that of their feather mites. However, we also explored the other τ values to obtain a more complete picture of the scaling of feather mite abundance. We used the *quantreg* R package (Koenker 2015), and assessed the slopes of the quantile regression models for different τ values from 0.05 to 0.95. Quantile regression analyses were also weighted by the sample size (log_10_ transformed) of each bird species.

Estimated mean λ for Q95 in the PGLS regressions explained above was 0.413 (95% CI: 0.077–0.749). Thus, a phylogenetic modelling approach to the quantile regression would require the phylogenetic scaling factor to be adjusted to λ<1. However, we were unaware of any tool able to perform such partial phylogenetic correction in a quantile regression analysis (see Jovani et al., 2016). Consequently, we present the results based on a non-phylogenetically corrected quantile regression and assume that phylogeny is unlikely to be a confounding factor.

Current information on the annual cycle of European feather mites indicates that their abundance peaks from winter until the onset of birds’ reproductive season (Blanco et al., 1997; Peet et al., 2022), when mites are transmitted from parents to offspring birds, causing a lowering of feather mite abundance (Mironov & Malyshev 2002; Doña et al., 2017). Thus, we tested the robustness of our conclusions by repeating all the analyses on feather mite abundance for the subset of birds captured from the beginning of October to the end of March (hereafter “winter”). This restriction reduced the sample size to 8,066 individual birds of 77 species.

## Results

### Predictions from Eq. 1 and Eq. 2

Microbial abundance on feathers was not correlated with bird species’ body size (Figure 2a, Table S1) which suggests that *σ*_h_would be 0. Moreover, bacteria and fungi are not produced by the host’s metabolism (in contrast to uropygial gland waxes). Thus, we removed −*γ*_h_ because it refers to a host mass-specific metabolic rate scaling. In this case, Eq. 1 predicts that feather mite abundance would only negatively scale with mite body size as follows (note that 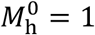)

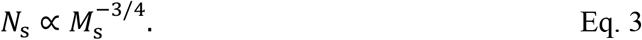

Uropygial gland size showed a strong allometric relationship with bird body size, with a scaling exponent of 0.902 (Figure 2b, Table S1). Therefore, if energy provided by the gland waxes of the hosts is the main constraint to feather mite infracommunities, Eq. 1 would predict that the maximum feather mite abundance would scale with bird and mite body size as follows

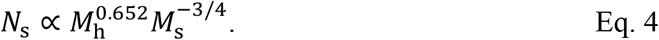

Thus, the scaling exponent of uropygial gland size on bird body size in Eq. 4 predicts a positive effect upon feather mite abundance because larger birds would provide more energy resources to mites. In contrast, the scaling exponent of feather mites’ body size is negative because (all else being equal) the higher energy consumption of larger mites would lead to lower abundances.

Wing area scaled with bird species body size to 0.676 power in accordance with the theoretical 2/3 scaling exponent for external host surfaces, while the number of barbs scaled with a slope of 0.264 (Figures 2c and 2d, Table S1). Thus, if feather mite infracommunities were limited by wing area, Eq. 2 would be

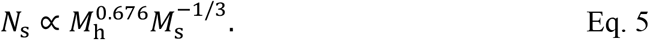

However, if the number of barbs is the relevant spatial constraint for feather mite infracommunities, Eq. 2 would read as

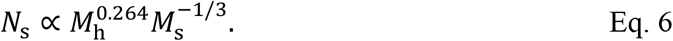

Thus, Eq. 5 and Eq. 6 show the predicted positive effect of bird body size upon feather mite abundance (larger birds provide more space to mites, depending on the bird’s body part relevant to the mites, i.e., wing area vs. barb amount), but that larger mites would attain a lower abundance (fewer mites would fit on a host of a given size).

### Predicted vs. empirical scaling rules

PGLS models showed a weak effect of feather mites’ body size on their abundance along all abundance quantiles (Table S2). For the few quantiles with slopes differing from 0, the slopes were all positive, thus strongly departing from the predicted negative slopes of –3/4 (Eqs. 3 and 4) and –1/3 (Eqs. 5 and 6) of mite abundance to mite body size (Table S2). In contrast, we found a positive correlation between bird species’ body size and the abundance of their feather mites (Figure 3, Table S2), holding from the Q45 to Q95, with empirical slopes in close agreement with the slopes predicted by Eq. 6 for the number of barbs (Figure 3a, Table S2). For the remaining host traits that might constrain mite abundances, empirical slopes departed from those predicted by the equations based on the scaling of microbial abundance (Eq. 3), uropygial gland size (Eq. 4), and wing area (Eq. 5; Figure 3a, Table S2).

**Figure 3:**
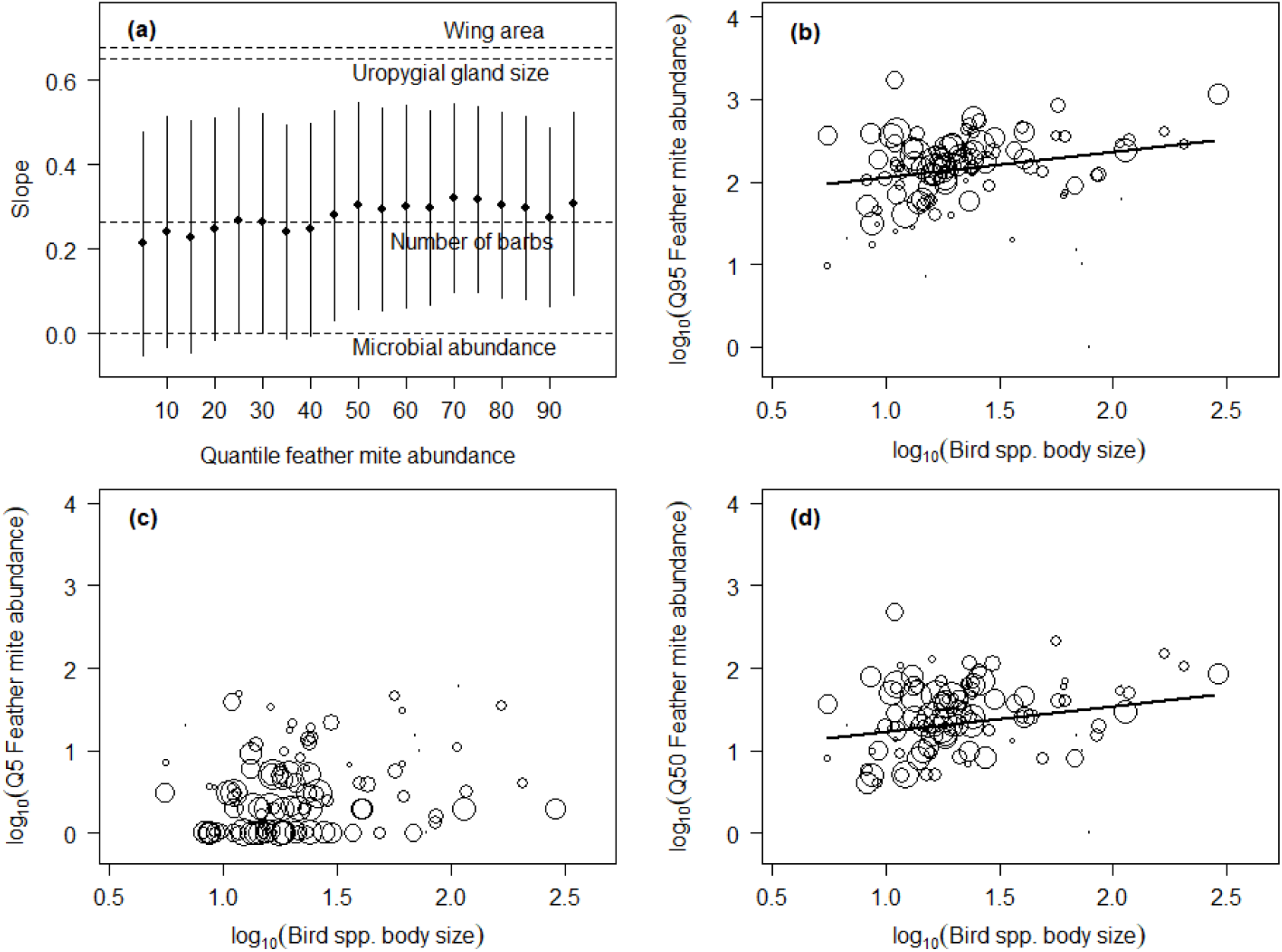
PGLS models for the relationship between 19 quantiles (from Q5 to Q95) of feather mite abundance in each bird species as dependent variable and log_10_(bird species body size) and log_10_(feather mite body size) as independent variables. (a) Slopes (±95% CI) for log_10_(bird species body size) are shown as dots and whiskers. Dashed lines show slope predictions according to Eq. 3 (microbial abundance), Eq. 4 (uropygial gland size), Eq. 5 (wing area), and Eq. 6 (number of barbs). (b) to (d) three examples of the relationship between bird body size (in g) and feather mite abundance at different quantiles (Q95, Q5, and Q50, respectively) from which slopes for plot (a) were obtained. Dot size is proportional to the log_10_(sample size) for each bird species. Only regression lines with slopes differing from 0 are shown.

Quantile regression analyses (including both bird and mite body size as independent variables) showed slopes for feather mite body size clearly departing from the predicted –1/3 and –3/4 by Eqs. 3 to 6 (particularly for higher τ values; Figure S1). However, we did found the expected positive relationship between Q95 feather mite abundance and bird species’ body size (Figure 4). Regression slopes for the highest τ values were close to the one predicted by the scaling equation that considers the number of barbs as a main spatial constraint (Eq. 6), but much larger than the slopes predicted for microbial abundance (Eq. 3), and much lower than those predicted by the scaling equations considering uropygial gland size (Eq. 4) or the wing area (Eq. 5) as the main energetic or spatial constraint, respectively (Figure 4).

**Figure 4:**
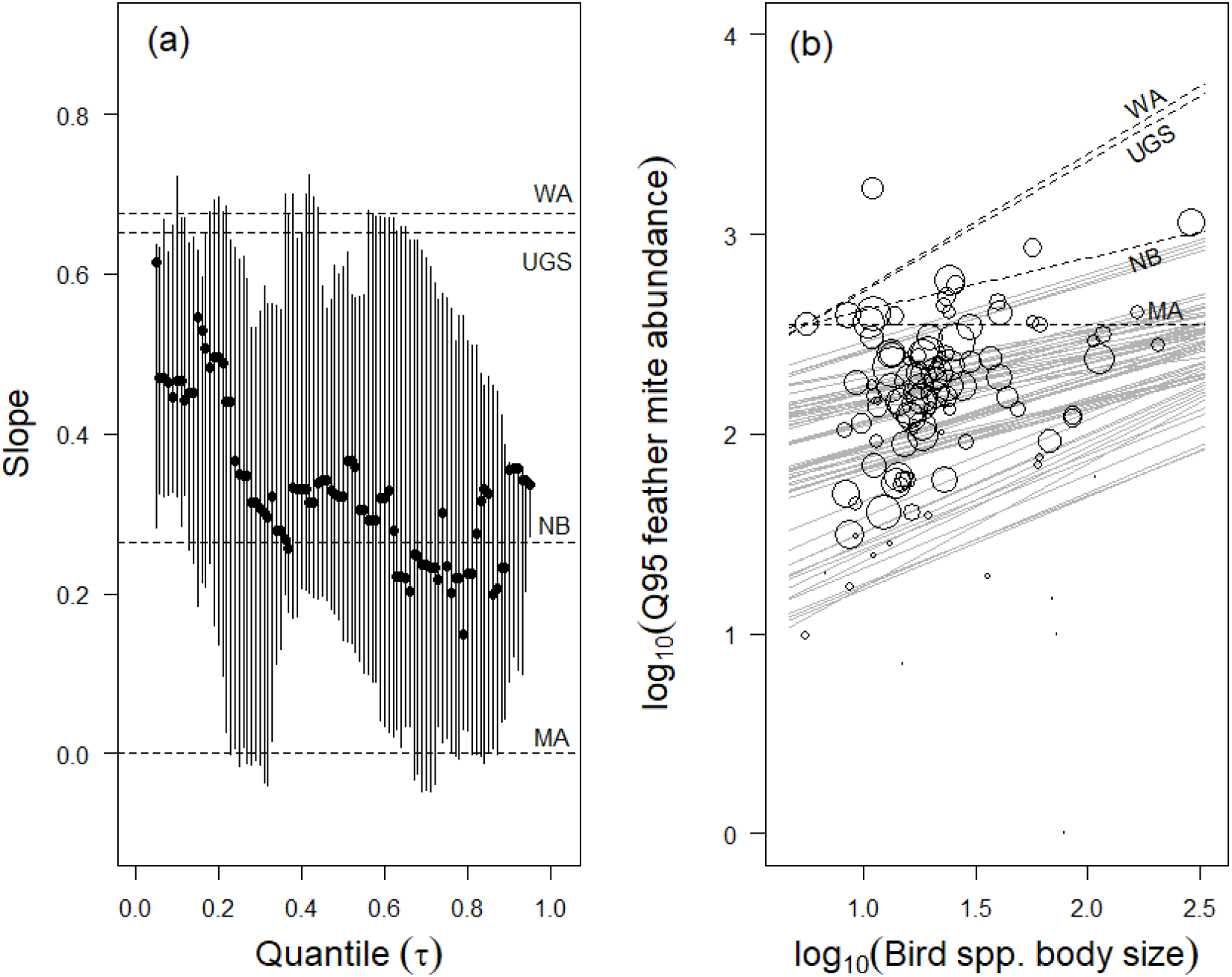
Multivariable quantile regression on log_10_(Q95 feather mite abundance) with log_10_(bird species body size) and log_10_ (feather mite body size) as independent variables. Dashed lines show slope predictions according to Eq. 3 (microbial abundance, MA), Eq. 4 (uropygial gland size, UGS), Eq. 5 (wing area, WA), and Eq. 6 (number of barbs, NB). (a) Bird species body size slopes (±95% CI) for each tau (τ) value. (b) Quantile regression of the allometry of Q95 feather mite abundance (continuous gray lines) and predicted slopes (black dashed lines). Dot size is proportional to the log_10_(sample size) for each bird species.

Feather mite’s body size was uncorrelated with hosts’ body size (Figure S2). Thus, overall, these results show that larger birds hold larger feather mite abundances, but this cannot be explained by feather mite size, i.e., larger birds do not carry larger numbers of smaller mites.

Dashed lines in Figure 4b (and Figure S3b) were drawn to cross the actual Q95 feather mite abundance (356 mites) for *Regulus ignicapilla*, the second smallest bird species in our sample (5.6 g). Thus, the dashed lines extrapolate the Q95 feather mite abundance expected for larger bird species, given the actual abundance of feather mites for smaller ones. Strikingly, this predicted that the largest bird species in our sample (*Pyrrhocorax pyrrhocorax*; 287.5 g) would have 5,098 and 4,635 Q95 feather mite abundance, according to the allometry of the wing area (Eq. 5) or the uropygial gland size (Eq. 4), respectively. However, the actual abundance was four times lower (1,155), and in close agreement with Eq. 6 (1,005) that involves the number of barbs. Also, the extrapolation according to Eq. 3 involving microbial abundance yielded a clear underestimation (356) of the actual abundance of mites. In summary, the rather flat slope of the quantile regression for the largest τ values (slope, 95% CI = 0.336, 0.272– 0.336) shows the strong ceiling that the number of barbs imposes on feather mites’ abundance, precluding larger birds from holding as many mites as expected based on other bird features.

When analysing only data from birds sampled in winter, the smaller sample size led to an increase in the uncertainty of the estimates, but similar qualitative results were found (see Supporting Information).

## Discussion

The carrying capacity of birds to support feather mite populations increases with bird species’ body size, with a scaling exponent close to that predicted by space (but not energy) constraints. Specifically, the empirical scaling we found fits closely the scaling exponents predicted by the equation involving the number of feather barbs, but not wing area as spatial constraint or microbial abundance or uropygial gland size as energetic constraints. The size of feather mites inhabiting bird species was not correlated with the abundance of mites or with the size of their hosts.

This space constraint seems to be in conflict with the fact that birds with many feather mites typically have large sections of each flight feather, or even entire feathers, devoid of feather mites (e.g. Jovani & Serrano 2004). However, feather mites show strong preferences for certain feathers and feather sections (e.g. Figure 1), and these preferences differ among feather mite species (Bridge 2003; Jovani & Serrano 2004; Mestre et al., 2011; Fernández-González et al., 2015; Stefan et al., 2015), feather mite life stages (Labrador et al., 2022), and according to environmental conditions (Wiles et al., 2000) or even to time of the day (Labrador et al., 2022). Therefore, our results, complemented with previous knowledge about the bird–feather mite system, show that feather mite populations are spatially limited, likely because of some negative density dependence acting well before the entire feather surfaces are fully occupied.

Our results simultaneously show a strong ceiling for the maximum feather mite abundance, and manifold differences in the abundance of feather mites among bird species with similar body sizes (note the logarithmic y-axis of Figures 3 and 4). For instance, in the Q95 abundance of feather mites of well-sampled bird species under 10 g there is an 8-fold difference from 47.9 mites per bird in *Phylloscopus collybita* to 389.5 mites in *Aegithalos caudatus*. Further comparative studies (as the one by Galván et al., 2008) are needed to understand which traits of birds and traits of feather mites are responsible for the large differences in feather mite abundances across bird species (Díaz-Real et al., 2014). Our results reject the role of uropygial gland size as an important constraint to feather mite populations and provide a new evidence (in addition to other studies, e.g. Pap et al., 2010; Doña et al., 2019) against preen waxes being important food resources for feather mites. Lastly, the role of bacteria and fungi as important food resources for feather mites would need further study because their potential limiting role should not be fully discarded given the small number of host species analyzed here in this regard (N = 21, Figure 2a).

Our results can be compared with those reported by Hechinger et al. (2019), who also studied the allometry of bird ectosymbionts’ abundance. While they found energetic constraints to be more relevant for arthropod ectosymbionts of birds, we have not found this energetic constraint. This disagreement may be because Hechinger et al. (2019) mainly studied non-passerine birds, and here we studied only passerines. Moreover, Hechinger et al. (2019) studied a more complete arthropod ectosymbiont community (lice and mites, including a few ticks), rather than focusing on a more taxonomically and ecologically restricted group as in our study (only feather mites). While there may be constraints shaping the whole community of ectosymbionts (thus supporting Hechinger et al.’s approach), it is also likely that different symbiont groups are constrained by different host traits, or by the same host traits but in different ways. For instance, unlike lice, blood-feeding mites and ticks, feather mites consume fungi, bacteria and other organic matter present on feathers’ surface, which are not produced by the host’s metabolism. Thus, this demands a different parameterization of the metabolic theory equations. Interestingly, Hechinger (2013) suggested that space constraints may be more relevant than energy in metabolically inactive symbiont stages that do not use the energy resources provided by their hosts (e.g. because they are trophically transmitted cyst stages waiting for an ultimate host to predate their current intermediate host). Our findings support this view as feather mites may not consume bird metabolic products, but organic material that they find on the feathers’ surface (Doña et al., 2019; Labrador et al., 2022). Thus, it is necessary to nurture the framework proposed by Hechinger (2013) and Hechinger et al. (2019) with more knowledge about the ecology and biology of the studied symbionts, and to integrate this with interspecific comparative analyses to understand the relevant processes regulating symbiont abundances and energy fluxes in host–symbiont systems.

The lack of correlation between the body size of the bird species studied here and the size of their feather mites goes against the Harrison’s Rule (Harrison 1915). This may be the result of feather mite species showing a complex co-evolutionary history with their hosts, with host-switching being as frequent as cospeciation (Doña et al., 2017, 2019). In other words, mites currently found on one bird species may have speciated on another host species (typically from the same genus or family). This may partly explain why the smallest (*Regulus regulus*; 5.6 g) and the largest (*Pyrrhocorax pyrrhocorax*; 287.5 g) bird species in our study have similarly sized mites (i.e., similar weighted mean size of their mite species): 3.82μg and 2.61 μg, respectively (Figure S2).

Besides the relevance of the number of barbs for mite abundance, the allometry of other host traits may also have interesting implications for our understanding of the entire symbiont community composed of all organisms living on bird feathers, the so-called pterosphere (*sensu* Labrador et al., 2021). For instance, we showed that feather mite abundance scaled with bird species’s body size with a much shallower slope than the wing area did (Figures 3 and 4, Table S2). Consequently, although absolute feather mite abundance increased with host body size, the maximum density of feather mites (i.e., Q95 feather mite abundance/cm^2^ wing area) decreased sharply with increasing bird species body size (PGLS: t = –3.083, df = 86, p = 0.003; Figures 5 and S5). This raises the question of (1) whether a lower density of feather mites in larger bird species implies a lower cleaning service provided by mites to their hosts; or (2) whether this lower density is the result of a potential competition between feather mites and feather lice, as numeric dominance of lice relative to mites has been observed in larger-bodied bird species (Hechinger et al., 2019).

**Figure 5:**
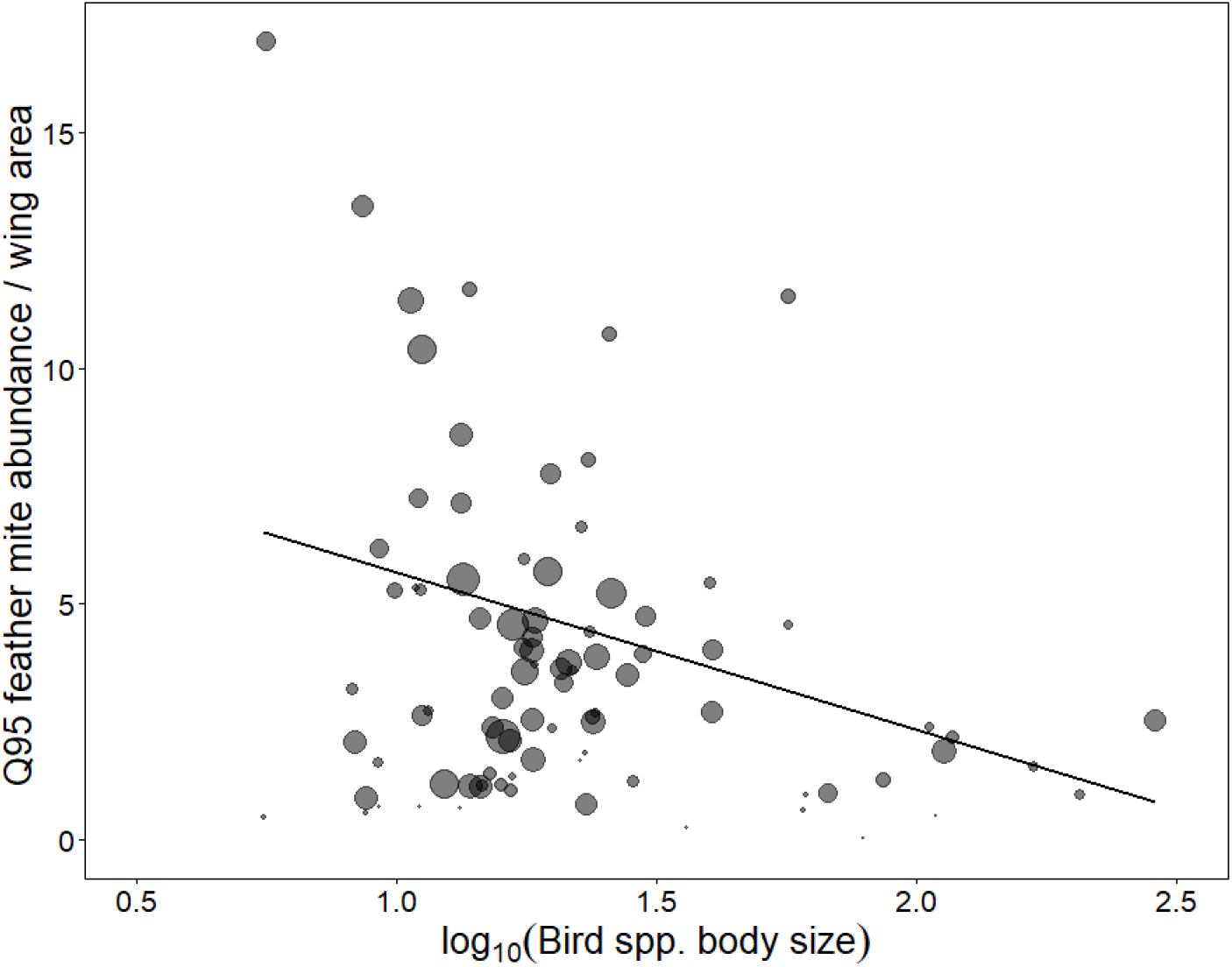
Relationship between log_10_(bird species body size) (in g) and the maximum density of their feather mites (Q95 feather mite abundance/cm^2^ of wing area). Dot size is proportional to the log_10_(sample size) for each bird species.

Overall, our study shows the potential of the theoretical and quantitative framework proposed by Hechinger (2013) using the metabolic theory of ecology to disentangle the mechanisms behind symbiont abundance across host species. It also shows the necessity to fully integrate the biology of the studied species to make accurate predictions on the factors limiting symbiont populations.

## Statement of authorship

ML, DS, RJ, MG conceptualised the study. All authosrs except for MG, LZG, and JD collected data. ML and RJ performed analyses with support from JD and LZG. ML and RJ wrote the first draft of the manuscript, and all authors contributed to revisions.

## Data accessibility statement

Data and code used in this study are available in the Dryad Digital Repository: https://datadryad.org/stash/share/LsvyoRzttwfkX1f-tUil5Fi2bJKNwJxeu48SX9VoIU0

## Acknowledgements

We are grateful to Brian S. Cade for his advice on quantile regression analyses. Manecorro Ringing Station (ICTS-RBD-CSIC) received economic support from the Junta de Andalucía. We thank Doñana National Park for the authorisations and support during the preparation of the ringing programmes. We thank all the ringers that help us with the measurements of the primaries at Manecorro Ringing Station. JD was supported by the European Commission (H2020-MSCA-IF-2019, INTROSYM:886532). SFG was supported by the Spanish Ministry of Economy and Competitiveness (CGL2010-15734/BOS). LZG was supported by a found from the National Research, Development, and Innovation Office – NKFIH (K-135841). OG was supported by the Spanish Ministry of Economy (Ref. CGL2014-56041-JIN). ML was supported by “la Caixa” Foundation (grant ID100010434). HP was supported by a Natural Sciences and Engineering Research Council of Canada (NSERC) Discovery Grant. JCS was supported by funds from the Ministry of Science and Innovation (CGL-2020 PID2020-114907GB-C21). CIV was supported by the Romanian Ministry of Research, Innovation and Digitization (CNCS - UEFISCDI; project no. PN-III-P1-1.1-TE-2021-0502).

