## Supporting Information for "Host space, not energy or symbiont size, constrains feather mite abundance across passerine bird species"

for Labrador et al.

***Feather mite's body size calculation***

For *Proctophyllodes* species, total feather mite length was measured from the pedipalp apex to the end of the hysterosomal lobes (see literature on feather mite morphology used in References section below). We also measured the length of the terminal appendages from scanned drawings of the same studies with Image J v1.53e (https://imagej.nih.gov/). Lastly, we added the two values to obtain a measure of the total length of each mite species.

For non-*Proctophyllodes* species, full body lengths measurements of females, including gnathosoma and terminal appendages (when relevant), were based on hard copies of published literature. A clear ruler was used to measure the length of the female's body and of the scale bar in mm, and the stated length of the scale bar (e.g. 200 µm), was used to convert the mm body length into µm. For 13 feather mite species (two *Joubertophyllodes* spp., five *Pteronyssoides* spp., five *Scutulanyssus* spp., and one *Sturnotrogus* sp.), the gnathosoma was not illustrated. In these cases, an illustration of a congeneric mite with the gnathosoma in dorsal view was used to estimate the proportion of the body made up of the gnathosoma, and that proportion was applied to the gnathosoma-free body lengths to estimate full body lengths with gnathosoma.

For *Trouessartia* species described before 1976, there were very few full-body illustrations available. Most of these *Trouessartia* species were redescribed by Santana

(1976). This author did not explicitly state how he measured ‘body length’, but provided only one full figure of a female of *T. poeopterae* [Gaud & Mouchet] (his Figure 3). Although there is no scale bar for this figure, Santana gave measurements of some prodorsal shield characters in the text. Using this stated length in µm of the prodorsal shield, we estimated the length in µm of the whole body, with and without gnathosoma, and also with and without the very small terminal lamellae on the terminal lobes. We then compared these length calculations to what Santana (1976) has in his text about the body length of female *T. poeopterae*. The measurement we made from the anterior margin of the prodorsal shield to the tips of the terminal lamellae was the closest match (just 7 µm difference) to body length described by Santana. So, we proceeded assuming that this was how he measured body length for all *Trouessartia* females. We then calculated the proportion of the total body length of *T. poeopterae* made up of the gnathosoma in dorsal view (8.2%) and applied that to all of the Santana (1976) body length measurements we used.

We utilized the equation made for Parasitidae Mesostigmata mites in Edwards (1967) to calculate the body size (in g) from their body length (and then divided it by 1000 to obtain the body size in µg).

$$Feather\ mite\ body\ size = \left(\frac{Feather\ mite\ body\ length}{3.95}\right)^3$$

We employed the Parasitidae equation rather than one for Astigmata because these feather mites are not globular/teardrop-shaped like the free-living Astigmata from which Edwards (1967) built the Astigmata equation. Instead, these feather mites are elongated and dorso-ventrally compressed like most Mesostigmata.

To our knowledge, the only measure of body size of a feather mite species is the one done by Gaede and Knülle (1987), who calculated that adult female *Proctophyllodes troncatus* [Mégnin] have an average body size of 3.49 µg. Our calculated value for this species was 2.64 μg, and thus potentially suggesting that our mite body size data may be slightly underestimated (note that this may differ for the other mite species). In any case, our study is interspecific and thus variance in body size across mite species (and not their absolute values) was the relevant feature in our present study.

### *Bird morphology data*

We acquired body masses of bird species from Dunning (2007). When both male and female body masses were reported for a species, the mean was calculated. We used European subspecies and populations when data for several subspecies or populations were reported. For the few species not included in Dunning (2007), data were obtained from the Birds of the World website (https://birdsoftheworld.org/bow/home).

Data on wing area were mainly obtained from Nudds et al. (2007), and the few species not reported there were retrieved from Pap et al. (2015) or Bruderer and Bolt (2001). Data on uropygial gland size were gathered from Vincze et al. (2013). We estimated the total number of barbs in primary feathers by multiplying the barb density of the P1 primary (i.e., the innermost; Pap et al., 2015) by the total (mean) length of primary feathers of each bird species. Barb density varies depending on the position of the feather, being higher in the distal ones (Pap et al., 2015). We used barb density data from P1 because it is the central wing feather, and therefore, potentially more representative, and because it was available for a larger number of species. The length of P1–P9 primary feathers (excluding the tiny outermost primary) was measured with a ruler (precision 0.5 mm) in 40,346 birds (sample size: mean = 917, min-max = 1–9,506 birds per species). These birds were captured and measured by JLA, AS, RR, BF, and other bird ringers from 1994 to 2015 at the Manecorro Ringing Station (Doñana National Park, SW Spain), during the autumn migration. Thus, primary feather lengths are representative of the Western European populations of the studied bird species. See Figure 2 in the main text for the number of bird species for each morphological variable.

***Results of analyses carried out considering only birds studied in winter***

PGLS regressions over quantiles showed similar estimates but with 95% CI crossing 0 (Figure 3 vs. S4; Table S2 vs. S3). Quantile regressions on the other hand showed the same estimates for higher τ values (the ones that show the ceiling imposed by the number of barbs on the abundance of mites; Figure 4 vs. S3). Regarding the effect of feather mite's body size on feather mite abundance, PGLS models (Table S2 vs. S3) and quantile regression analyses (Figure S1 vs. S6) led to equal conclusions.

**Table S1.** PGLS model coefficients of the relationship between potential energy and space constraints against bird species body size (in g) (see also Figure 2 in main text). CI.l and CI.u show lower and upper values of the 95% confidence interval for slope, respectively.

| *Variable* | *Intercept* | *Slope* | *CI.l* | *CI.u* |
|---|---|---|---|---|
| Fungi and bacteria | -1.618 | -0.0648 | -0.573 | 0.444 |
| Uropygial gland size | 0.882 | 0.9019 | 0.792 | 1.011 |
| Wing area | -3.188 | 0.6761 | 0.624 | 0.728 |
| Number of barbs | 2.839 | 0.2638 | 0.189 | 0.339 |

**Table S2.** Coefficients of the PGLSs models between $\log_{10}$(bird species' body size) and $\log_{10}$(feather mite's body size) as independent variables and 19 quantiles (from Q5 to Q95) of feather mite abundance in each bird species.

| | | *Bird body mass* | | | *Feather mite body mass* | | |
|---|---|---|---|---|---|---|---|
| *Quantile* | *Intercept* | *Slope* | *95%CI* | | *Slope* | *95%CI* | |
| Q5 | 0.092 | 0.214 | -0.051 | 0.478 | -0.059 | -0.560 | 0.443 |
| Q10 | 0.203 | 0.241 | -0.033 | 0.514 | 0.093 | -0.427 | 0.613 |
| Q15 | 0.363 | 0.229 | -0.044 | 0.503 | 0.062 | -0.459 | 0.582 |
| Q20 | 0.487 | 0.248 | -0.014 | 0.510 | 0.077 | -0.422 | 0.576 |
| Q25 | 0.573 | 0.268 | 0.002 | 0.534 | 0.085 | -0.422 | 0.592 |
| Q30 | 0.663 | 0.263 | 0.006 | 0.520 | 0.129 | -0.362 | 0.619 |
| Q35 | 0.779 | 0.241 | -0.013 | 0.496 | 0.149 | -0.334 | 0.633 |
| Q40 | 0.830 | 0.247 | -0.006 | 0.499 | 0.196 | -0.283 | 0.674 |
| Q45 | 0.861 | 0.280 | 0.030 | 0.529 | 0.201 | -0.269 | 0.672 |
| Q50 | 0.930 | 0.304 | 0.059 | 0.549 | 0.171 | -0.292 | 0.634 |
| Q55 | 1.018 | 0.295 | 0.054 | 0.536 | 0.166 | -0.286 | 0.619 |
| Q60 | 1.078 | 0.302 | 0.063 | 0.541 | 0.185 | -0.262 | 0.633 |
| Q65 | 1.178 | 0.298 | 0.068 | 0.529 | 0.172 | -0.259 | 0.602 |
| Q70 | 1.219 | 0.320 | 0.097 | 0.543 | 0.225 | -0.190 | 0.641 |
| Q75 | 1.300 | 0.317 | 0.097 | 0.537 | 0.264 | -0.143 | 0.670 |
| Q80 | 1.385 | 0.304 | 0.084 | 0.523 | 0.313 | -0.091 | 0.718 |
| Q85 | 1.501 | 0.299 | 0.082 | 0.516 | 0.344 | -0.053 | 0.740 |
| Q90 | 1.618 | 0.275 | 0.064 | 0.487 | 0.431 | 0.045 | 0.817 |
| Q95 | 1.751 | 0.308 | 0.091 | 0.525 | 0.424 | 0.028 | 0.821 |

**Table S3.** As Table S2, but only analyzing data from birds sampled in winter.

| | | *Bird body mass* | | | *Feather mite body mass* | | |
|---|---|---|---|---|---|---|---|
| *Quantile* | *Intercept* | *Slope* | *95%CI* | | *Slope* | *95%CI* | |
| Q5 | 0.371 | 0.078 | -0.285 | 0.441 | -0.020 | -0.708 | 0.668 |
| Q10 | 0.636 | 0.026 | -0.343 | 0.395 | 0.077 | -0.619 | 0.773 |
| Q15 | 0.792 | 0.011 | -0.365 | 0.387 | 0.109 | -0.601 | 0.819 |
| Q20 | 0.846 | 0.079 | -0.285 | 0.443 | 0.137 | -0.549 | 0.823 |
| Q25 | 0.892 | 0.110 | -0.265 | 0.485 | 0.172 | -0.535 | 0.878 |
| Q30 | 1.019 | 0.097 | -0.261 | 0.454 | 0.154 | -0.521 | 0.830 |
| Q35 | 1.041 | 0.119 | -0.234 | 0.471 | 0.222 | -0.444 | 0.888 |
| Q40 | 1.146 | 0.130 | -0.209 | 0.470 | 0.183 | -0.456 | 0.823 |
| Q45 | 1.186 | 0.145 | -0.190 | 0.479 | 0.207 | -0.424 | 0.837 |
| Q50 | 1.210 | 0.180 | -0.146 | 0.506 | 0.214 | -0.399 | 0.828 |
| Q55 | 1.193 | 0.228 | -0.089 | 0.545 | 0.263 | -0.335 | 0.861 |
| Q60 | 1.220 | 0.248 | -0.063 | 0.560 | 0.302 | -0.285 | 0.890 |
| Q65 | 1.304 | 0.242 | -0.066 | 0.551 | 0.303 | -0.280 | 0.886 |
| Q70 | 1.370 | 0.248 | -0.046 | 0.543 | 0.310 | -0.247 | 0.867 |
| Q75 | 1.437 | 0.264 | -0.023 | 0.551 | 0.273 | -0.269 | 0.814 |
| Q80 | 1.509 | 0.272 | -0.012 | 0.555 | 0.292 | -0.246 | 0.829 |
| Q85 | 1.593 | 0.267 | -0.016 | 0.551 | 0.320 | -0.217 | 0.858 |
| Q90 | 1.725 | 0.252 | -0.030 | 0.533 | 0.358 | -0.178 | 0.894 |
| Q95 | 1.861 | 0.236 | -0.056 | 0.528 | 0.436 | -0.119 | 0.992 |

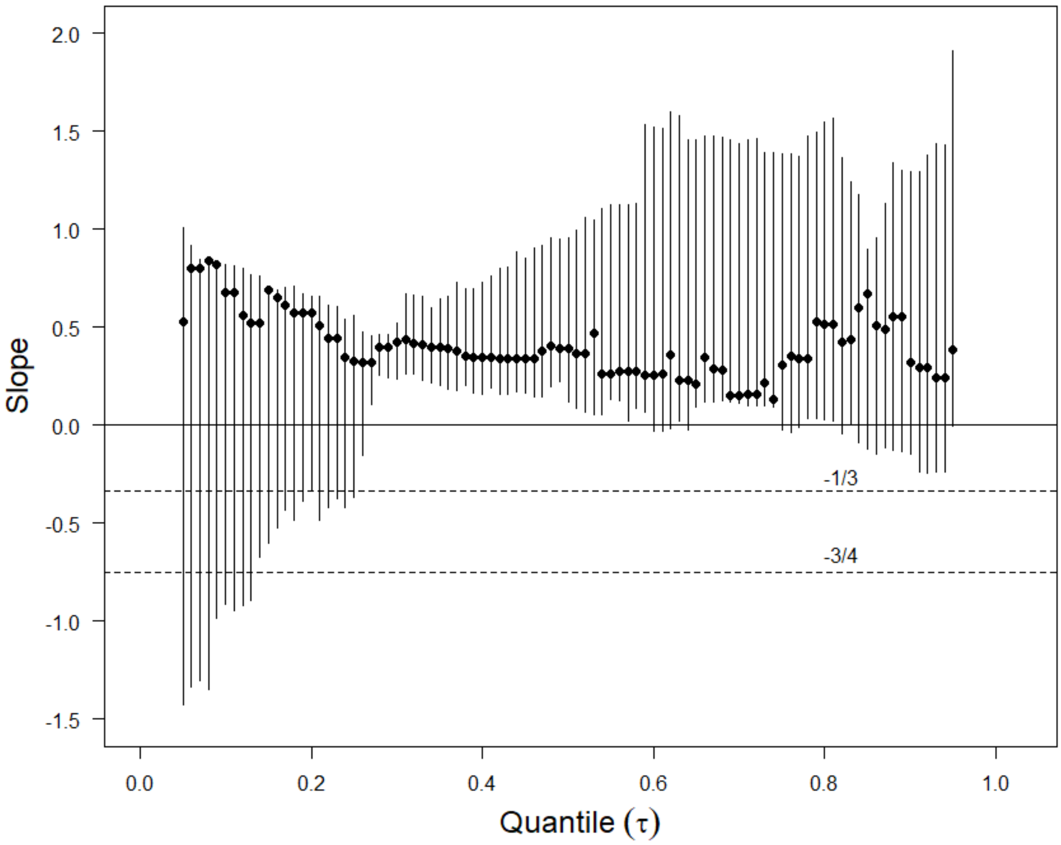


**Figure S1:** Multivariable quantile regression on $\log_{10}$(Q95 feather mite abundance) with $\log_{10}$(bird species body size) and $\log_{10}$(feather mite's body size) as independent variables. Dots (±95% CI) show feather mite body size slopes for each tau (τ) value. Dashed lines show slope predictions according to Eqs. 3 and 4 (slope = –3/4) for energy and Eqs. 5 and 6 (slope = –1/3) for space constraints on feather mite abundance.

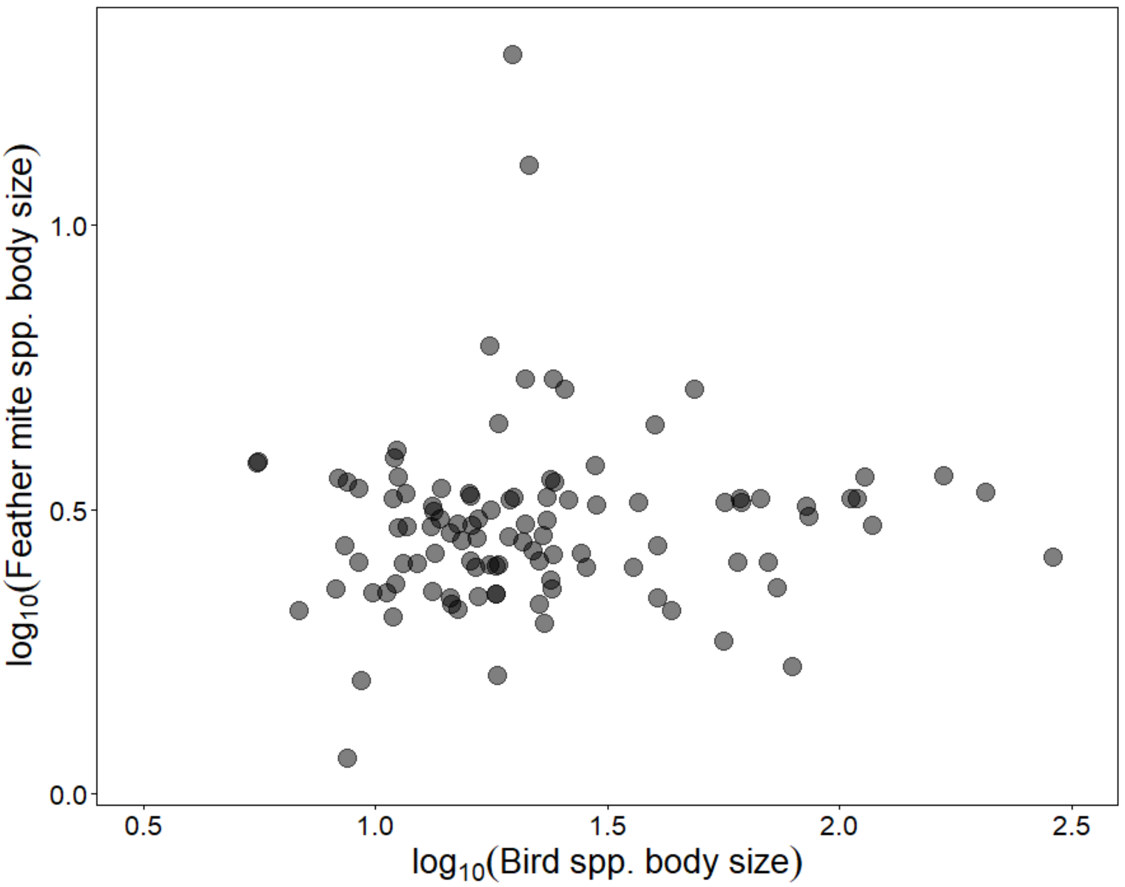


**Figure S2:** Relationship between the weighted mean of feather mite species body size (in µg) and the body size (in g) of their bird host species (PGLS t = 0.660, df = 104, p = 0.511).

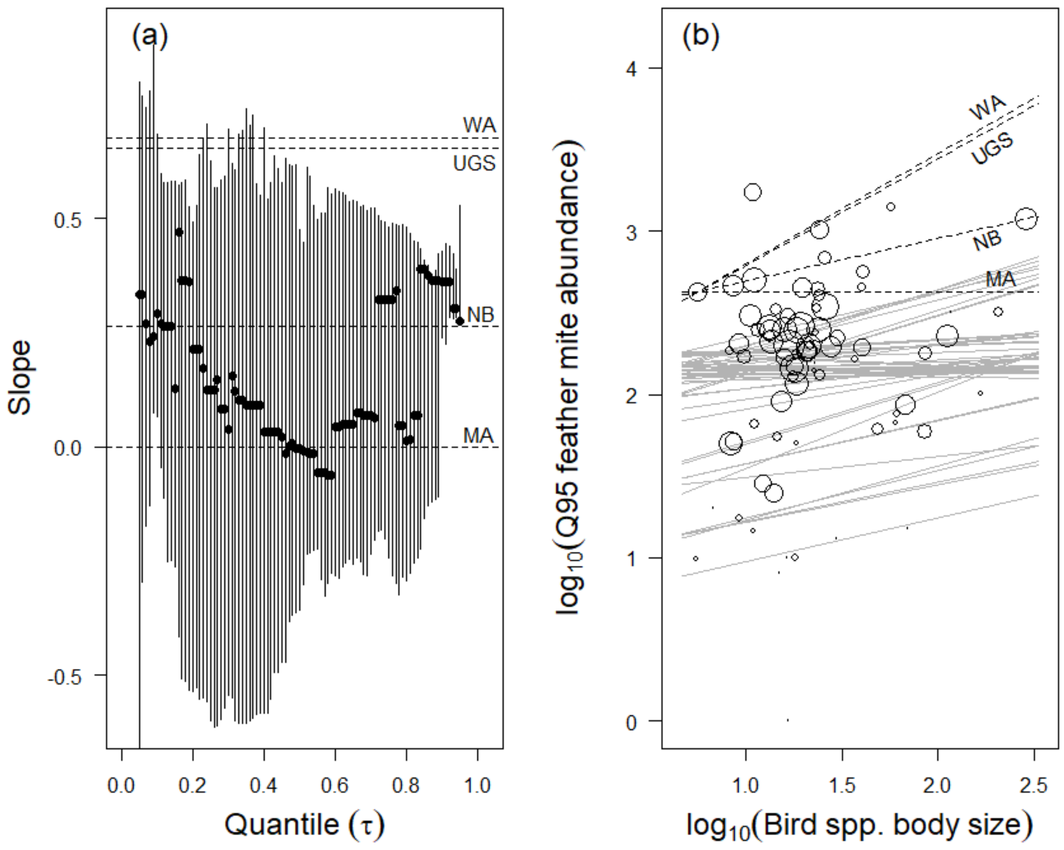


**Figure S3:** As in Figure 4 in the main text, but only data from birds sampled in winter are used. Multivariable quantile regression on $\log_{10}$(Q95 feather mite abundance) with $\log_{10}$(bird species body size) and $\log_{10}$(feather mite body size) as independent variables. Dashed lines show slope predictions according to Eq. 3 (microbial abundance, MA), Eq. 4 (uropygial gland size, UGS), Eq. 5 (wing area, WA), and Eq. 6 (number of barbs, NB). (a) Bird species' body size slopes (±95% CI) for each tau (τ) value. (b) Quantile regression of the allometry of Q95 feather mite abundance (continuous gray lines) and predicted slopes (black dashed lines). Dot size is proportional to the $\log_{10}$(sample size) for each bird species.

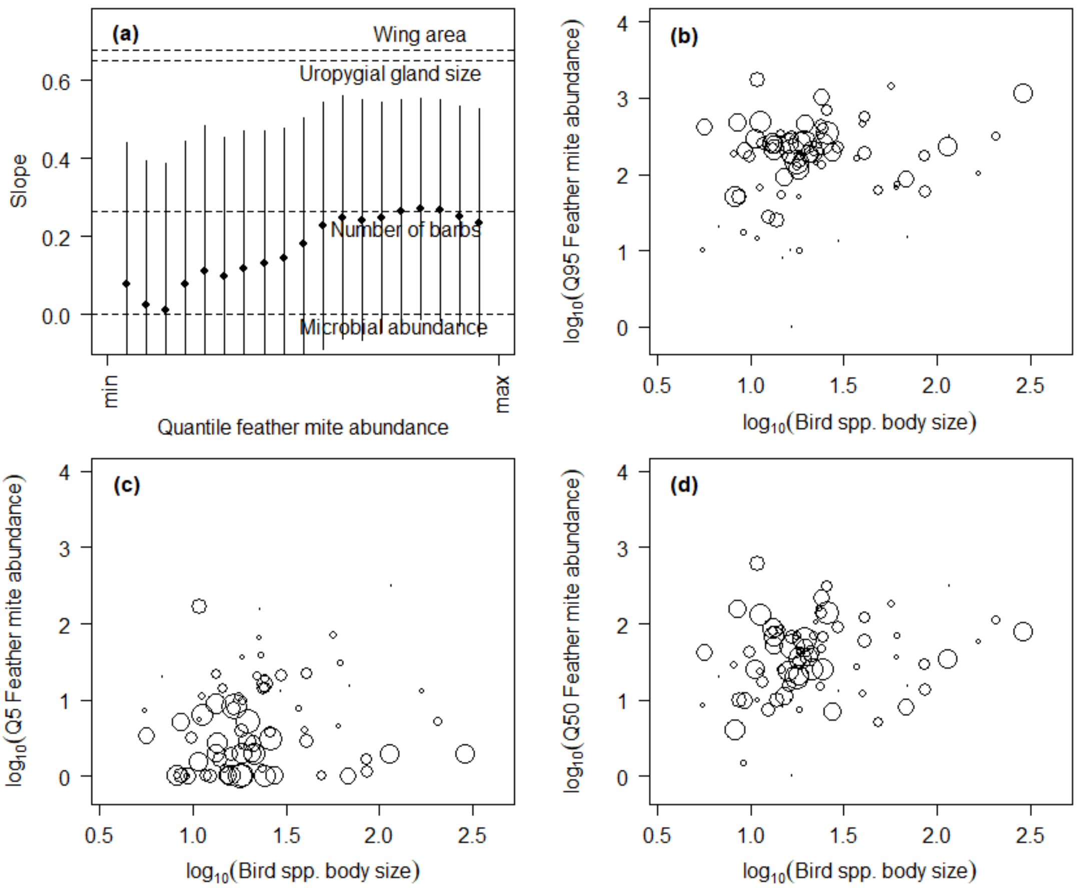


**Figure S4:** As Figure 3 in the main text, but only data from birds sampled in winter. (a) PGLS models between 19 quantiles (from Q5 to Q95) of feather mite abundance in each bird species as dependent variable and $\log_{10}$(bird species' body size) and $\log_{10}$(feather mite's body size) as independent variables. Slopes (±95% CI) for $\log_{10}$(bird species' body size) are shown. Dashed lines show slope predictions according to Eq. 3 (microbial abundance), Eq. 4 (uropygial gland size), Eq. 5 (wing area), and Eq. 6 (number of barbs). (b) to (d) three examples of the relationship between bird body size (in g) and feather mite abundance from which slopes in (a) were obtained. Dot size is proportional to the $\log_{10}$(sample size) for each bird species.

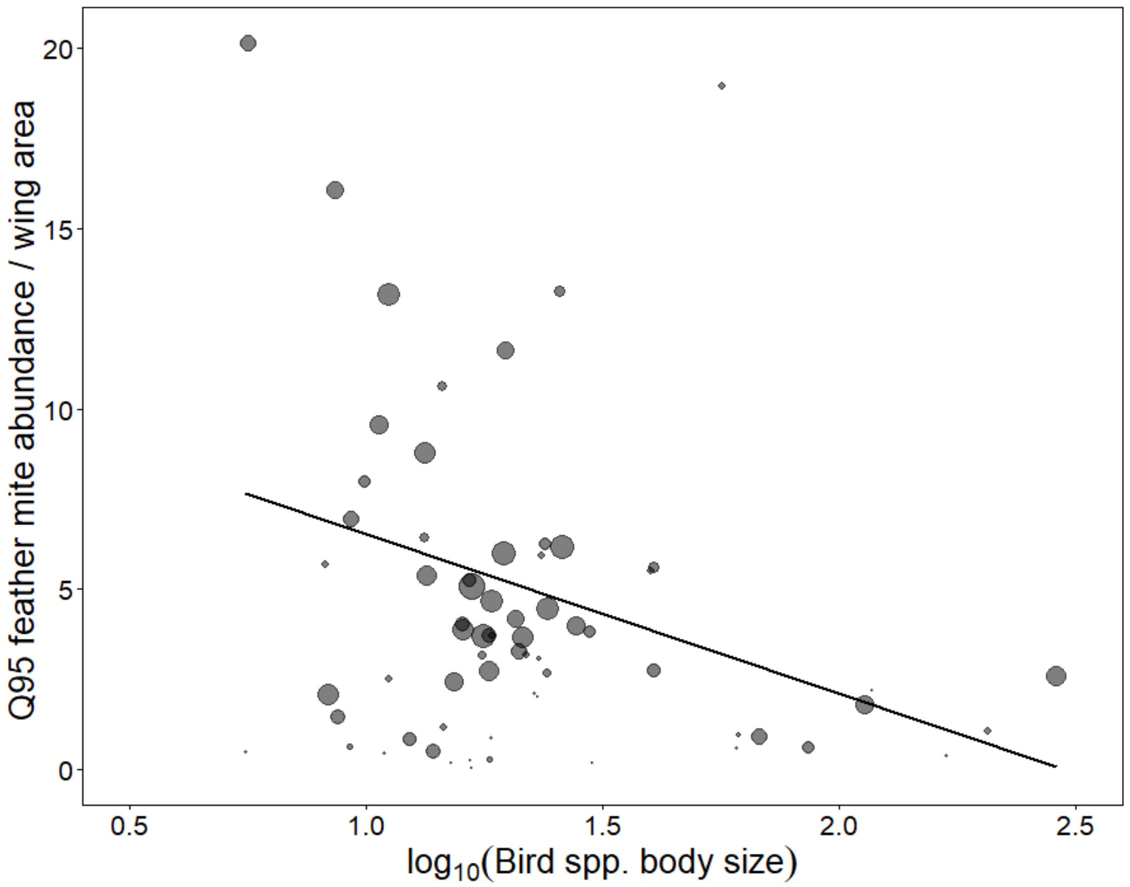


**Figure S5:** As Figure 5 in the main text, but only data from birds sampled in winter were used. PGLS between $\log_{10}$(bird species body size) (in g) and their maximum density of feather mites (Q95 feather mite abundance/$cm^2$ of wing area) ($t = -3.004$, $df = 65$, $p = 0.004$). Dot size is proportional to the $\log_{10}$(sample size) for each bird species.

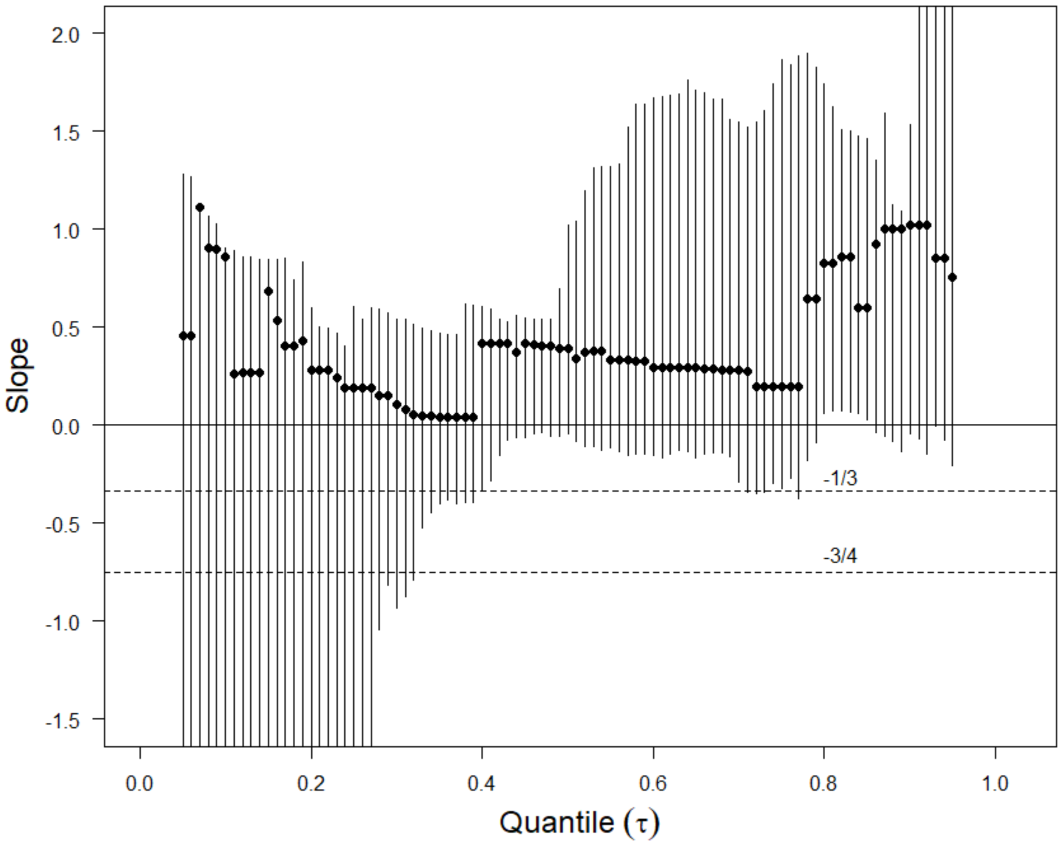


**Figure S6:** As Figure S1, but only analysing birds sampled in winter. Multivariable quantile regression on $\log_{10}$(Q95 feather mite abundance) with $\log_{10}$(bird species body size) and $\log_{10}$ (feather mite body size) as independent variables. Dots (±95% CI) show feather mite body size slopes for each tau (τ) value. Dashed lines show slope predictions according to Eqs. 3 and 4 (slope = –3/4) for energy and Eqs. 5 and 6 (slope = –1/3) for space constraints on feather mite abundance.
